# Glutamate and γ-aminobutyric acid differentially modulate glymphatic clearance of amyloid β through pulsation- and aquaporin-4 dependent mechanisms

**DOI:** 10.1101/2020.01.31.928481

**Authors:** Cheng Wu, Yi-wei Feng, Qun Zhang, Feng-yin Liang, Yue Lan, Zhong Pei, Guang-qing Xu

## Abstract

The glymphatic system contributes to a large proportion of brain waste clearance, including removal of amyloid β (Aβ). We have demonstrated that glutamate and γ-aminobutyric acid (GABA) influence glymphatic clearance through distinct mechanisms whereby GABA exerts modulatory effects in an aquaporin-4 (AQP4)-dependent manner while the actions of glutamate are pulsation-dependent. The efficacy of GABA and glutamate in alleviating Aβ in APP-PS1 and Angiotensin-II (Ang-II) induced hypertension mouse models was further evaluated. Notably, increasing GABA or inhibiting glutamate levels led to reduced binding of Aβ to pre-labeled plaques to similar extents in APP-PS1 mice while GABA appeared more efficient in Aβ clearance in hypertensive animals than the glutamate inhibitor. Our findings support the modulation of neurotransmitters that influence the glymphatic pathway via distinct mechanisms as a potentially effective therapeutic strategy for clearance of Aβ deposits from the brain.

## Introduction

β-Amyloid (Aβ) accumulation is one of the pathological hallmarks of Alzheimer’s disease (AD). Both genetic and environmental factors are involved in AD development. Earlier studies have provided substantial evidence supporting an important role of environmental factors, such as hypertension, in the development of sporadic AD^1^. Cerebral blood vessels provide major Aβ clearance routes from the brain. Hypertension disrupts the normal function of cerebral blood vessels, promoting vascular Aβ deposition, which, in turn, impairs vessel function, forming a vicious circle and eventually leads to cognitive dysfunction^2, 3^.

In the CNS, Aβ is released from neurons into brain interstitial fluid (ISF). Recently, the glymphatic system and dural lymphatic vessels have been proposed as major extracellular disposal routes of Aβ^4–7^.

Neurotransmitters play an important role in the pathogenesis of AD. Aβ is reported to disrupt synaptic transmission of different neurotransmitters^8–11^, leading to cognitive decline. Several pre-clinical and clinical studies have demonstrated that modulation of neurotransmitter systems by memantine (NMDA receptor antagonist) and etazolate (GABA-A receptor agonist) can effectively alleviate accumulation of Aβ^12, 13^. Interestingly, neurotransmitters are also involved in modulation of glymphatic clearance. For example, previous studies have shown that noradrenaline mediates glymphatic transport in that convective exchange is increased in the presence of adrenergic antagonists, resulting in faster clearance of Aβ^14^. Glutamate and GABA are major stimulatory and inhibitory neurotransmitters in the CNS, respectively. However, their potential roles in the glymphatic system are yet to be established. Aquaporin-4 (AQP4) and arterial pulsation are two main driving forces in the glymphatic pathway^4, 15^. The GABA-A receptor co-localizes with AQP4 in brain tissue^16, 17^. In addition, glutamate has been shown to influence vascular smooth muscle cell function, which is associated with pulsation^18^. In the current study, we tested the hypothesis that glutamate and GABA play important roles in glymphatic pathway clearance through modulation of pulsation or AQP4. Improper function of these neurotransmitters may differentially contribute to clearance failure of Aβ.

## METHODS

### Animals

C57BL/6J male mice (6–8 weeks, 12–16 weeks and 8 months) were provided by the Sun Yat-sen University Medical Experimental Animal Center (Guangzhou, China). AQP4-deleted male mice (AQP4^−/−^) 6–8 weeks of age were a gift from the Jiangsu Key Laboratory of Neurodegeneration (Nanjing Medical University, Nanjing, China). Male APPswe/PS1dE9 (APP/PS1) mice 7–8 months old were purchased from Guangdong Medical Experimental Animal Center. AQP4^−/−^ and APP-PS1 mice were both backcrossed onto a C57BL/6J background. All animals were kept in temperature- and humidity-controlled rooms under a 12 h/12 h light/dark cycle. Sample size was selected according to previously published reports. Experimental procedures were performed in accordance with the guidelines imposed by Sun Yat-sen University Committee on the Care and Use of Animals.

### Reagents and Antibodies

FITC-dextran 70 kDa (Sigma, FD4, USA) was used to trace CSF movement, and FITC-dextran 2000 kDa (Sigma, FD2000S, USA) and Rhodamine B-dextran 70 kDa (Sigma, R9379, USA) to label vasculature. All reagents were dissolved in artificial CSF (ACSF) at a concentration of 1%. Neurotransmitter agonists/antagonists, including glutamate (Sigma, G3291, USA), NMDA receptor antagonist, 2-amino-5-phosphonovaleric acid (APV) (Sigma, A8054, USA), AMPA/kainate receptor antagonist 6-Cyano-7-nitroquinoxaline-2, 3-dione (CNQX) (Sigma, C127, USA), GABA (Sigma, A2129, USA), GABA-A receptor antagonist and Bicuculline (Bic) (Sigma, 285269, USA), were dissolved in CSF tracer before use at a concentration of 0.2 mM. Evans Blue (Sigma, E2129, USA) dissolved in saline was used to label lymph node drainage. Angiotensin-II (Bechem, H-1705, Torrance, USA) was employed to induce hypertension. Fluorescent-labeled Aβ peptides (Anaspec, 60492-01, Fremont, USA) and FSB (Millipore, 344101, Darmstadt, Germany) were used to examine Aβ binding. To label amyloid plaques, antibodies specific for Aβ 1-42 (Biolegend, 805501, San Diego, USA) and Aβ 1-40 (Biolegend, 805401, USA) were employed. Antibodies for Collagen-I (Co-I) (Abcam, ab34710, Hong Kong, UK) and α-smooth muscle actin (SMA) (Boster, BM0002, Wuhan, China) were used to test vascular structure changes and those for AQP4 (Alomone Labs, 300-314, Jerusalem, Israel) and GFAP (Millipore, 2642205, USA) to detect aquaporin and astrocytes, respectively. All secondary antibodies were purchased from Cell Signaling Technology (4408, 4409, 4412, 4413, Carlsbad, USA).

### Intra-cisternal and interstitial tracer injection

Mice were anesthetized with an intraperitoneal injection of 1% pentobarbital (50 mg/kg) and positioned in a stereotaxic frame (RWD Life Science company, Shenzhen, China). A microsyringe (BASi, West Lafayette, USA) was inserted into the cisterna magna (i.c.v.), and 10 μl CSF tracer injected at a speed of 0.2 μl/min. For interstitial injection, micro glass pipettes filled with 1 μl CSF tracer was connected to a microinjection system (BASi, USA). Under a two-photon microscope, the pipette was inserted into the cortex under 200 μm and the solution infused at a velocity of 0.2μl/min

### *In vivo* two-photon imaging of glymphatic pathway clearance

A 2×2mm^2^ cranial window was prepared over the right parietal cortex (2 mm caudal from bregma, 1.7 mm lateral from the midline) with a micro-drill and a metal plate glued at the edge of the cranial window^16, 17^. The mouse was fixed on the stage of two-photon microscope (Leica, DM6000, Wetzlar, Germany). ACSF was perfused during the whole surgical procedure to keep the cranial window moist. Throughout the experiment, body temperatures were kept constant at 36.8°C with a feedback-controlled heating pad (RWD Life Science Company, Shenzhen, China). To visualize vasculature, 0.2 ml Rhodamine B-dextran 70 kDa was injected intravenously before imaging. A Leica NA 0.95 together with a 25× magnification water-immersion objective was used at an excitation wavelength of 800 nm. The cerebral vasculature was initially imaged with 512 × 512-pixel frames from the surface to a depth of 200 μm with 2 μm z-steps. Tracer movement was detected with dual channels. Imaging panels 100 μm below the cortical surface were selected for analysis of tracer movement into the paravascular space with Leica Lite software. For paravascular and interstitial movement, circular regions of interest (ROI) 25 pixels in diameter were centered on the surrounding penetrating arterioles. To define tracer movement into brain tissue, the distribution of CSF tracers in three-dimensional (3D) vectorized reconstruction was analyzed. Mean pixel intensities within these ROIs and 3D reconstructions were measured at 5 min intervals. For interstitial clearance, mean fluorescence of CSF tracers was measured at 15 min intervals.

### Evans blue injection and quantification

With careful separation of muscle and anadesma in the back, the cisterna magna was exposed. In total, 5 μl of 10% Evans blue (Sigma, E2129) was slowly injected into the cisterna magna at a rate of about 1 μl/min over 5 min. After 30 min, deep cervical lymph nodes (dcLNs) were dissected for assessment of Evans blue content. The intensity of Evans blue was measured using confocal microscopy, and the same capture parameters maintained during intensity detection.

### Pulsation measurement

Pulsation measurements were conducted as previous experiment described^15^. X–T line scan technology was used to measure pulsation. Scans of 2000 ms (0.5 ms per line) were acquired orthogonal to the vessel axis in different vessel types, including surface arteries, surface veins, penetrating arteries and penetrating veins. Vessel fluorescence was extracted from X-T plots and plotted in relation to time using Graphpad prism 6.0. Pulsatility was calculated as standard deviation of the percentage of mean values.

### Unilateral internal carotid artery ligation

Unilateral internal carotid artery ligation was performed as previous experiment described^15^. The right common, internal, and external carotid arteries were surgically isolated. The internal carotid was ligated by 5-0 silk suture.

### Murine models of hypertension

Hypertension mouse models were generated by Angiotensin-II (1000 ng/kg/min) infusion using osmotic pumps (DURECT, Alzet model 2004), which were implanted subcutaneously in 12–16 weeks-old C57Bl/6 mice for 28 days. Assessment of Ang-II amounts was performed according to previous reports^19^. Blood pressure was measured with the tail cuff method every week. Mouse models without hypertension were discounted.

### *In vivo* two-photon time-lapse imaging of Aβ binding

Mice received FSB injection (I.P. 7.5 mg/kg) two days prior to the experiment. Anesthesia and craniotomy were performed in keeping with previous protocols. Microglass pipettes filled with 1 μl Aβ-555 peptide solution were connected to a microinjection system (BASi, USA). Aβ40-555 were used among hypertension groups and Aβ42 555 were used among APP/PS1 groups. Under a two-photon microscope, the pipette was inserted into the cortex and the solution infused at a velocity of 0.2 μl/min. *In vivo* two-photon imaging of the binding process was conducted about 200 μm away from the pipette tip with a 20× water immersion objective. Aβ40/42-555 and FSB were excited at 920 nm and 800 nm, respectively. The recording time was 10 min with 30 s per frame. For administration of the GABA, APV and CNQX, drugs were dissolved in a CSF at a final concentration of 10 μM in 10 μl and injected into the cisternal at the velocity of 1μl/min.

### Immunofluorescence

Immunofluorescence analysis was conducted on 10 μm paraformaldehyde (PFA)-fixed frozen sections. Slices were blocked for 1 h at room temperature with normal goat serum and 0.3% Triton and incubated overnight with primary antibody at 4°C, followed by secondary antibody for 1 h at room temperature. Slices were stained with DAPI (F6057, Sigma, USA) and immunofluorescence images observed under a confocal microscope (Leica, DM6000, Wetzlar, Germany).

### Statistical analysis

Data were analyzed with SPSS Statistics 20 and GraphPad Prism 6.0 and presented as mean ± s.e.m or mean ± S.D. Different treatment groups were evaluated using one-way ANOVA with LSD test or two-way ANOVA with Tukey’s test for multiple comparisons to determine differences among individual groups. The unpaired t-test was used when comparing two individual groups. A probability of p<0.05 was indicative of significant differences between groups. Regardless of the method used, the results were equivalent in magnitude and statistically significant. Data from all statistical analyses are presented in Supplementary Table 1.

## RESULTS

### GABA promotes while glutamate suppresses glymphatic clearance

To explore the roles of GABA and glutamate in glymphatic pathway clearance, specific agonists/antagonists for these neurotransmitters dissolved in CSF tracer were injected into the cisterna magna (Fig. 1e). Cerebral vasculature was labeled with 70 kDa Rhodamine B, which allows visualization of arteries based on blood flow direction and morphology.

**Fig. 1.**
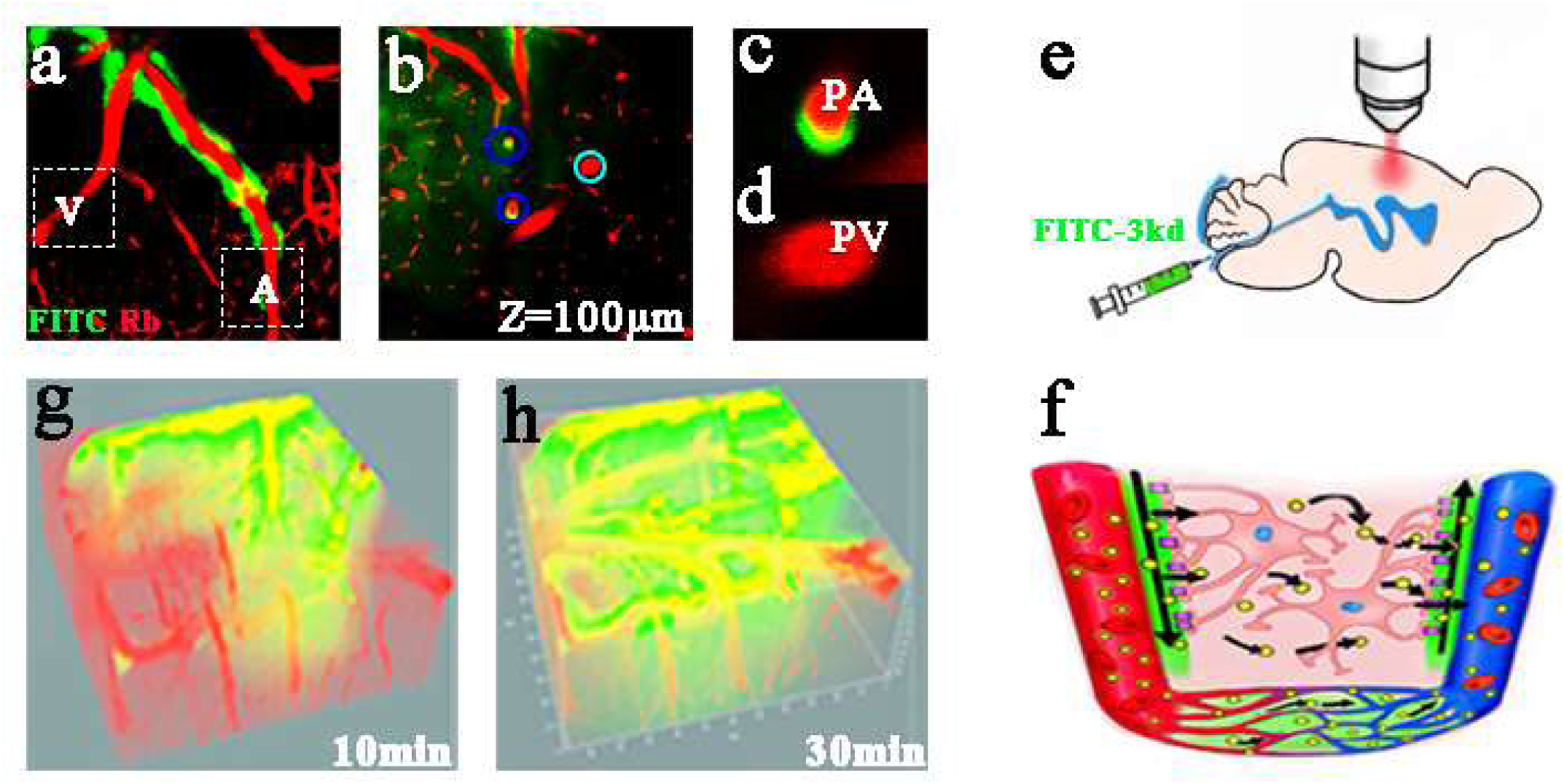
*In vivo* two-photon imaging of CSF tracer clearance through the glymphatic system. **(a–d)** *In vivo* imaging of CSF tracer movement along the surface **(a)** and penetrating arteries **(b)**. Cerebral vasculature was visualized with Rhodamine B (red). CSF tracer (green) moved along the paravascular space of arteries but not veins (A: surface artery; V: surface vein; PA: penetrating artery; PV: penetrating vein; Dark blue circles, arterioles; light blue circles, veins). Magnification of penetrating vessels **(c)** and **(d)**. **(e)** Schematic diagram of the imaging setup and intra-cisternal injection of the CSF tracer. **(f)** Schematic depiction of the glymphatic system. In this brain-wide pathway, CSF enters the brain along para-arterial routes and is cleared along paravenous routes. **(g–h)** 3D reconstruction of distribution of the CSF tracer in brain at 10 and 30 min. The tracer moved rapidly into the parenchyma over time.

Measurements were taken 5 min after infusion when the fluorescent tracer could be stably visualized. The tracer rapidly entered the cortex through the paravascular spaces surrounding the surface along penetrating arteries but was absent in venules at the beginning (Fig. 1a–d). To quantitatively evaluate the movement of para-arterial CSF tracer, mean fluorescence intensity (ROI) was measured at 100 μm below the cortical surface over 30 min with intervals of 5 min (Fig. 1b). During the measurements, mean fluorescence intensity in the sham group increased constantly up to ∼300% at 30 min (Fig. 2a-c). A large proportion of para-arterial CSF tracer gradually fluxed into this space. Interestingly, glutamate significantly inhibited paravascular penetration (Glutamate vs. vehicle, two-way ANOVA, for interaction factor, *P*<0.001) while both the GABA_-A_ receptor agonist and antagonist had no effect on paravascular movement (GABA vs. vehicle, two-way ANOVA, for interaction factor, *P*=0.6027; bicuculline vs. vehicle, two-way ANOVA, for interaction factor, *P*=0.2211) (Fig 2b). The neurotransmitter glutamate has two major ionotropic receptors (NMDA and AMPA/kainate receptors), which are widespread in the cerebral cortex and other brain regions. Notably, upon separate blockage of these two receptors, APV (NMDA receptor antagonist) significantly accelerated paravascular movement (APV vs. vehicle, two-way ANOVA, for interaction factor, *P*<0.001) whereas CNQX (AMPA/kainate receptor antagonist) had no effect (CNQX vs. vehicle, two-way ANOVA, for interaction factor, *P*=0.9303). Since water and small solutes freely enter the brain interstitium from paravascular spaces via bulk flow, we additionally analyzed tracer influx into the surrounding interstitium via 3D vectorized reconstruction of tracer distribution in the parenchyma (Fig. 2b–c). The mean fluorescence intensity in the sham group increased to ∼3.0-fold at 30 min, indicating influx of tracer from the paravascular space. As expected, glutamate reduced CSF tracer penetration into the interstitium (Glutamate vs. vehicle, two-way ANOVA, for interaction factor, P<0.001) (Fig. 2d). Unexpectedly, APV and CNQX robustly accelerated tracer influx into parenchyma (APV vs. vehicle, two-way ANOVA, for interaction factor, *P*=0.1346; CNQX vs. vehicle, two-way ANOVA, for interaction factor, *P*<0.001) (Fig. 2d). Moreover, GABA induced marked enhancement whereas bicuculline (GABA-A receptor antagonist) led to significant inhibition of influx (GABA vs. vehicle, two-way ANOVA, for interaction factor, *P*<0.05; bicuculline vs. vehicle, two-way ANOVA, for interaction factor, *P*<0.001) (Fig. 2e). Therefore, our results suggest that glutamate slows paravascular movement though the NMDA receptor and inhibits tracer influx into the parenchyma though the AMPA/kainate receptor. GABA, which did not influence paravascular movement, accelerated tracer influx into the parenchyma.

**Fig. 2.**
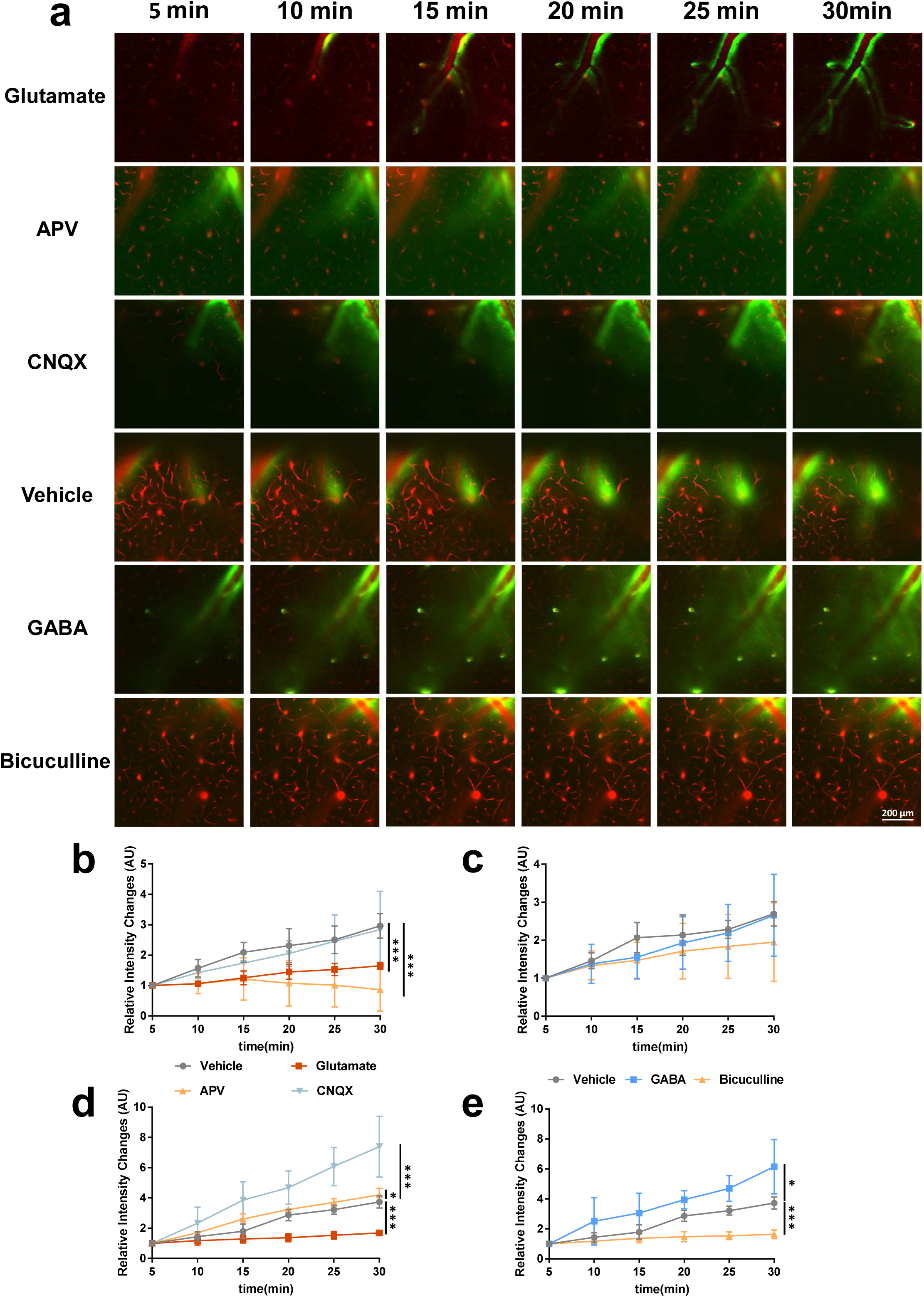
GABA promotes while glutamate suppresses glymphatic clearance. **(a)** *In vivo* two-photon imaging of tracer clearance through the para-vascular glymphatic pathway after intra-cisternal injection of GABA/glutamate and the respective antagonists. **(b, c)** Quantification of CSF tracer influx into the surrounding parenchyma via 3D reconstruction and paravascular CSF tracer clearance at 100 μm below the cortical surface **(d, e)**. Glutamate strongly inhibited whereas APV accelerated paravascular movement. No significant changes in paravascular movement were observed in mice receiving GABA, bicuculline and CNQX. Glutamate inhibited penetration of the tracer into the interstitium. APV did not influence the dynamic pattern of tracer penetration while CNQX significantly promoted tracer influx. GABA promoted whereas bicuculline inhibited tracer influx to a significant extent (n=6 mice per group). Present and following dgata are presented as means ± S.D. \**P*<0.05, \*\**P*<0.01 and \*\*\**P*<0.001.

In the glymphatic pathway, CSF enters the brain interstitium though paravascular movement and interstitial fluid in parenchyma is subsequently cleared from paravenous routes. To further explore the role of neurotransmitters in modulating interstitial fluid clearance, CSF tracers were injected directly into brain parenchyma 150-200 μm below the skull surface and two-photon imaging of tracer clearance performed over 60 minutes with intervals of 15 minutes (Fig. 3a). The mean fluorescence intensity in the region of interest (ROI) at 200 μm below the cortical surface was measured to quantify interstitial fluid clearance. Following injection, mean fluorescence intensity in the vehicle group constantly decreased and was reduced by ∼60% at 60 minutes (Fig. 3b, c). Notably, glutamate strongly inhibited while CNQX significantly accelerated interstitial fluid clearance (glutamate vs. vehicle, two-way ANOVA, for interaction factor, *P*<0.001; CNQX vs. vehicle, two-way ANOVA, for interaction factor, *P*<0.001). However, APV did not promote interstitial fluid clearance (APV vs. vehicle, two-way ANOVA, for interaction factor, *P*=0.6069). In contrast, GABA greatly promoted whereas bicuculline robustly inhibited interstitial fluid clearance (GABA vs. vehicle, two-way ANOVA, for interaction factor, *P*<0.001; bicuculline vs. vehicle, two-way ANOVA, for interaction factor, *P*<0.01) (Fig. 3b, c). Based on these findings, we concluded that glutamate impairs interstitial fluid clearance though the AMPA/kainate receptor and GABA facilitated drainage.

**Fig. 3.**
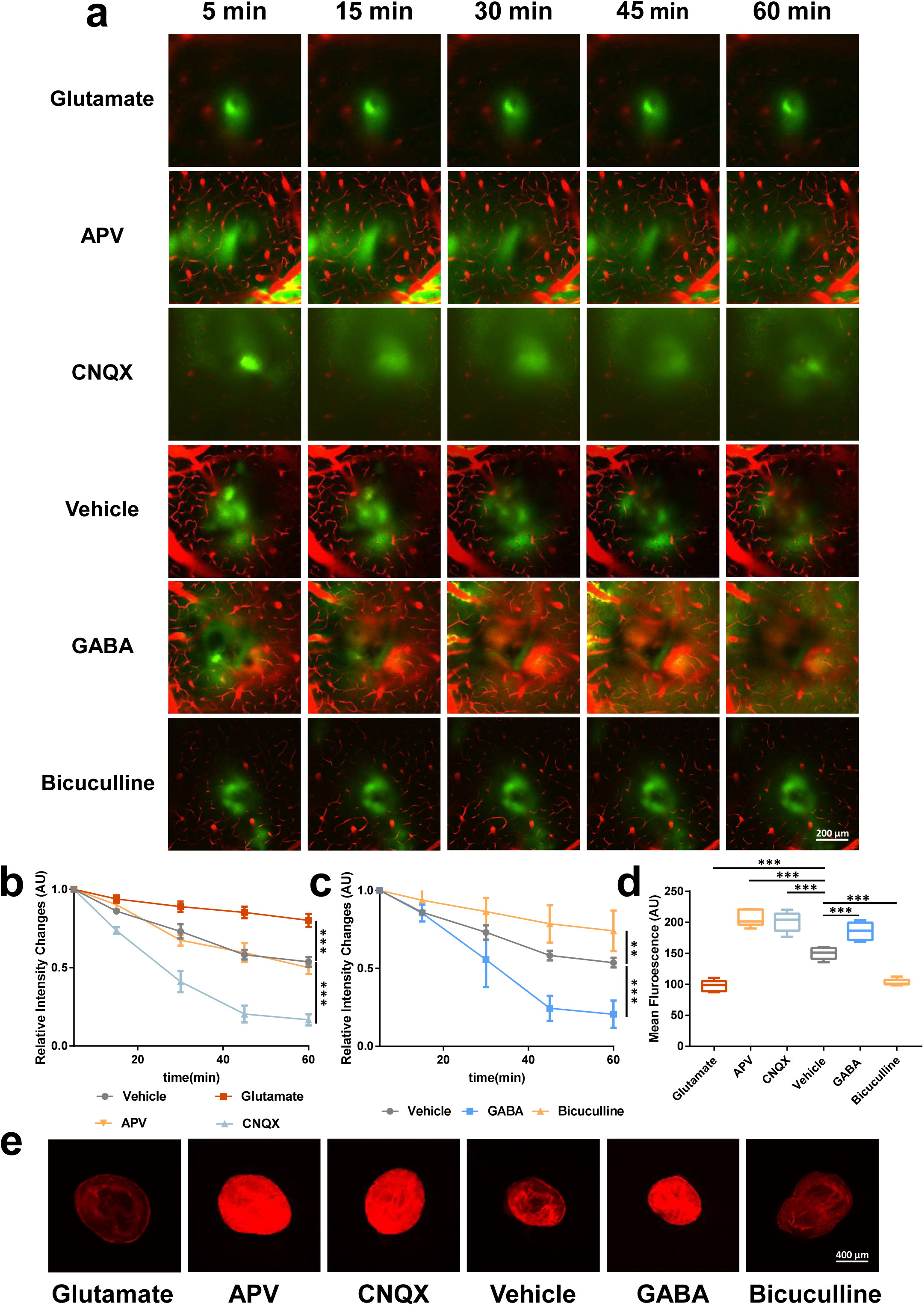
Glutamate inhibits while GABA accelerates ISF clearance. **(a)** *In vivo* two-photon imaging of tracer clearance in the interstitium after injection of tracer containing glutamate/GABA agonist/antagonist. **(b, c)** Quantification of mean tracer fluorescent changes in mice receiving glutamate/GABA agonist/antagonist. Glutamate significantly inhibited whereas CNQX accelerated tracer clearance. ISD clearance was promoted by GABA and conversely inhibited by bicuculline. **(e)** Representative fluorescent images of Evans blue staining indicative of drainage into dcLNs after injection of GABA/glutamate and the respective antagonists. dcLNs in each group are magnified by 50×. **(d)** Quantification of dcLN fluorescence in each group. Glutamate significantly inhibited whereas CNQX and APV promoted drainage. Moreover, drainage was enhanced by GABA and inhibited by bicuculline (n=6 mice per group).

Interstitial solutes are ultimately cleared into peripheral lymph nodes outside the brain parenchyma. Accordingly, we injected Evans blue i.c.v. and examined for the presence of the dye in deep cervical lymph nodes (dcLNs) in animals receiving different treatments. Consistent with previous findings, Evans blue was detected in dcLNs 30 min after injection. Mean fluorescence of Evans blue in lymph nodes was analyzed for each group (Fig. 3e). Unexpectedly, Evans blue staining was barely detectable in dcLNs of animals receiving glutamate (Fig. 3b) but evident in lymph nodes of animals receiving APV and CNQX, respectively (Glutamate vs. vehicle, *t*=9.976, *P*<0.001; APV vs. vehicle, *t*=8.737, *P*<0.001; CNQX vs. vehicle, *t*=6.811, *P*<0.001) (Fig. 3d), suggesting that a glutamate-related clearance mechanism is involved in this drainage route of cerebrospinal fluid (CSF). Consistently, GABA accelerated whereas bicuculline inhibited ultimate drainage of the tracer dye (GABA vs. vehicle, *t*=5.27, *P*<0.001; bicuculline vs. vehicle, *t*=10.85, *P*<0.001) (Fig. 3e). Our findings suggest that glutamate and GABA modulate drainage of interstitial fluid from the parenchyma via distinct routes. Based on the collective findings, we propose that glutamate and GABA play opposite roles in CSF tracer movement. Specifically, glutamate inhibits paravascular movement though the NMDA receptor and facilitates interstitial fluid clearance though the AMPA/kainate receptor while GABA does not influence paravascular movement but accelerates interstitial fluid drainage.

### Glutamate-mediated inhibition of paravascular movement though the NMDA receptor is pulsation-dependent

Paravascular movement of CSF is reported to driven by arterial pulsation. Our previous experiments showed that glutamate inhibits paravascular movement though the NMDA receptor. Accordingly, we proposed that glutamate-mediated inhibition of paravascular movement was pulsation-dependent.

To further examine this hypothesis, two-photon line scanning to visualize pulsation was performed. Vascular pulsatility was measured at different levels of the cerebrovascular tree, specifically, surface arteries, penetrating arteries, ascending veins and surface veins (Fig. 4a). In the vehicle group, penetrating arteries and ascending veins showed significantly higher pulsatility, compared with surface arteries and veins (Fig. 4b, c). Upon administration of glutamate and its antagonist (i.c.v.), glutamate strongly inhibited whereas APV promoted pulsatility among surface and penetrating arteries (Among surface arteries, Glutamate vs. vehicle, *t*=6.498, *P*<0.001; APV vs. vehicle, *t*=5.613, *P*<0.001; among penetrating arteries, Glutamate vs. vehicle, *t*=6.437, *P*<0.001; APV vs. vehicle, *t*=13.400, *P*<0.001) (Fig. 4d). However, CNQX, GABA, and bicuculline had no significant influence on pulsatility (Fig. 4e). We concluded from this experiment that glutamate reduced pulsatility though the NMDA receptor while GABA did not influence pulsatility.

**Fig. 4.**
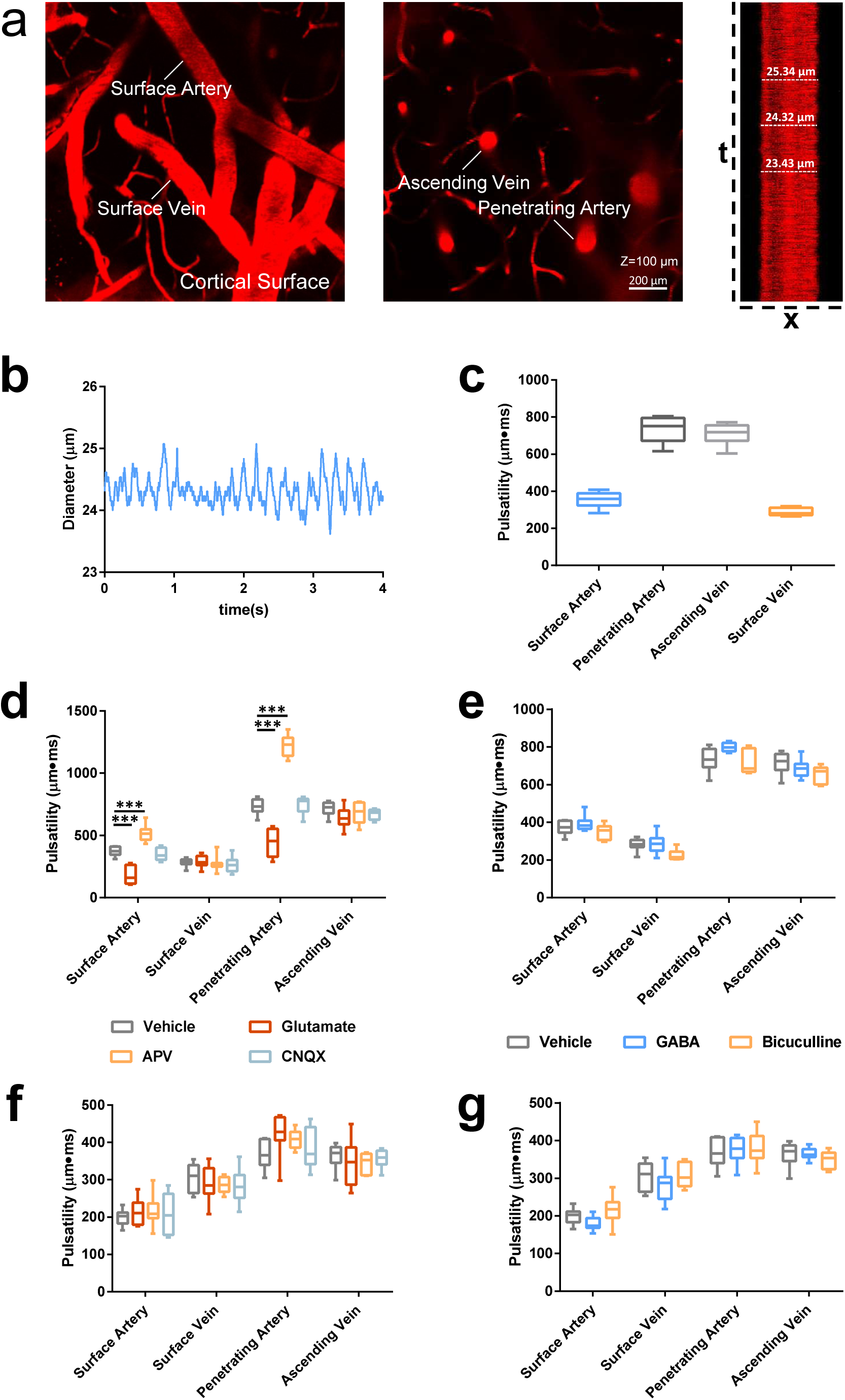
Measurement of vascular pulsatility in mouse cortex. **(a)** Cortical surface arteries and veins and penetrating arteries and veins. X–T line scans (red lines) were generated orthogonal to the vessel axis. **(b)** Vascular pulsatility was defined as absolute changes in fluorescence values over 4000 ms with intervals of 0.5 ms. **(c)** Vascular pulsatility in cortical surface arteries (SA), penetrating arteries (PA), ascending veins (AV) and surface veins (SV). Pulsatility was greatest in penetrating arteries and veins, compared with surface vessels. **(d-e)** Analysis of pulsatility in different groups. Glutamate strongly inhibited whereas APV significantly accelerated pulsatility among surface and penetrating arteries. GABA and bicuculline did not affect pulsatility. **(f-g)** Analysis of pulsatility in different groups after deep cerebral artery ligation. Artery ligation abolished pulsatility changes by glutamate and APV (n=8-9 vessels per group).

Based on pulsatility measurements, we further reduced cerebral arterial pulsatility by unilateral ligation of the internal carotid artery and injected the CSF tracer (i.c.v.), which dissolves glutamate and the GABA agonist/antagonist. After arterial ligation, we observed significantly reduced pulsatility in the vehicle group (Fig. 5b). Moreover, administration of glutamate and GABA agonist/antagonist failed to induce significant changes in pulsatility (Fig. 5c). Next, we quantified paravascular tracer movememt 100 μm below the cortex surface as described previously. The vehicle group showed markedly slower paravascular movement after arterial ligation (Fig. 5a). Moreover, glutamate and APV failed to either inhibit or facilitate paravascular movement (Glutamate vs. vehicle, two-way ANOVA, for interaction factor, *P*=0.3392; APV vs. vehicle, two-way ANOVA, for interaction factor, *P*=0.2709) (Fig. 5b). Meanwhile, administration of CNQX, GABA, and bicuculline did not lead to significant changes in paravascular movement, compared with the vehicle group (CNQX vs. vehicle, two-way ANOVA, for interaction factor, *P*=0.4799; GABA vs. vehicle, two-way ANOVA, for interaction factor, *P*=0.8563; bicuculline vs. vehicle, two-way ANOVA, for interaction factor, *P*=0.9873) (Fig. 5c). The mean fluorescence intensity in parenchyma was additionally measured. The vehicle group showed markedly slower tracer accumulation in parenchyma (Fig. 5d, e). However, glutamate and APV groups displayed almost no CSF tracer penetration into parenchyma (Glutamate vs. vehicle, two-way ANOVA, for interaction factor, *P*=0.9991; APV vs. vehicle, two-way ANOVA, for interaction factor, *P*=0.9693) (Fig. 5d). Interestingly, GABA and CNQX remarkably facilitated tracer influx (GABA vs. vehicle, two-way ANOVA, for interaction factor, *P*<0.001; CNQX vs. vehicle, two-way ANOVA, for interaction factor, *P*<0.05) (Fig. 5e).

**Fig. 5.**
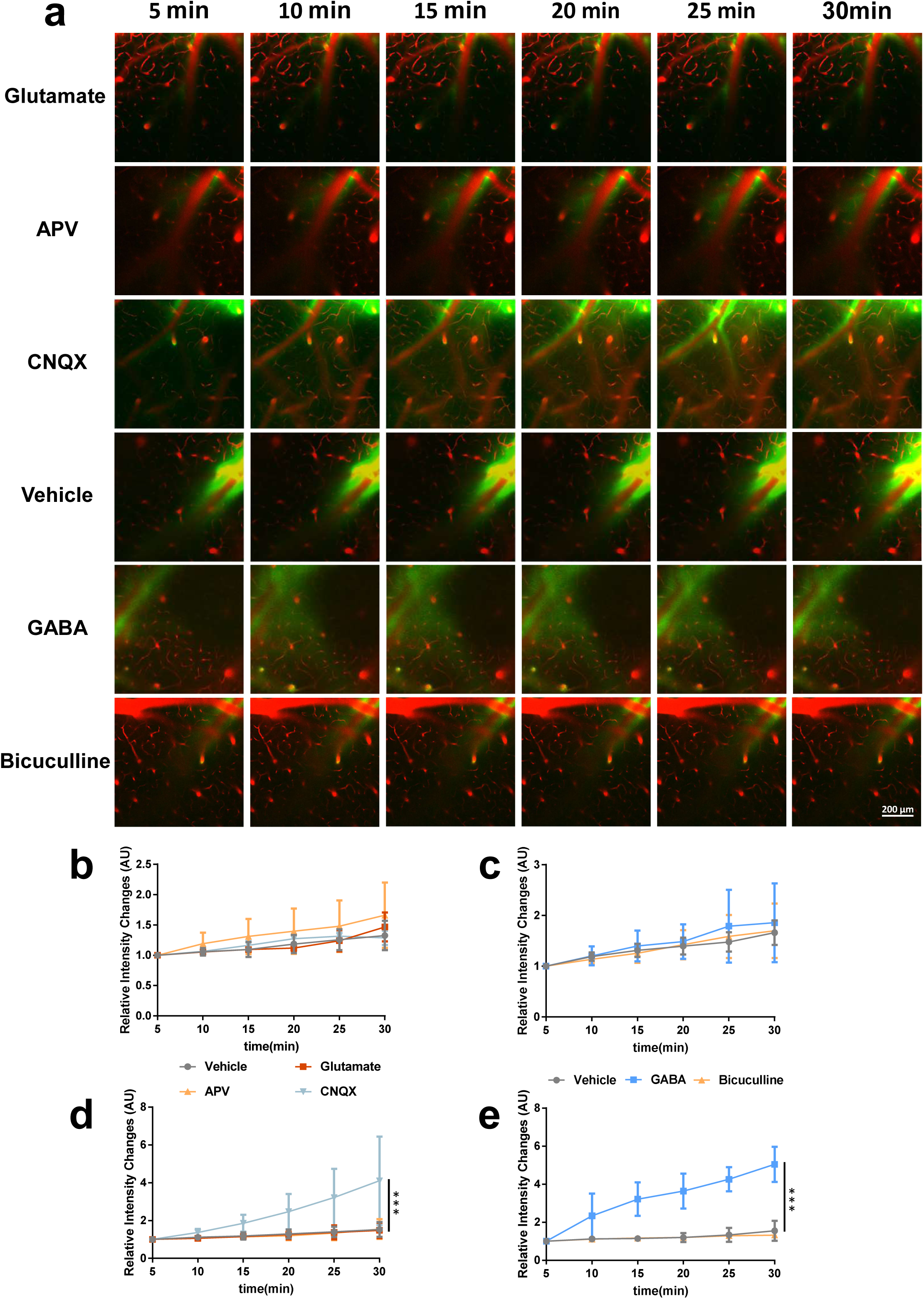
Glutamate inhibits paravascular movement though the NMDA receptor, which is pulsation-dependent. **(a)** *In vivo* two-photon imaging of tracer clearance through the para-vascular glymphatic pathway after intra-cisternal injection of GABA/glutamate and the respective antagonists. Deep cervical artery was unilaterally ligated during imaging. **(b, c)** Quantification of CSF tracer influx into the surrounding parenchyma via 3D reconstruction and paravascular CSF tracer clearance at 100 μm below the cortical surface **(d, e)**. After ligation, no significant differences were observed in mice receiving glutamate and GABA agonists/antagonists with regard to paravascular movement. CNQX and GABA significantly accelerated tracer penetration into the interstitium (n=5-6 mice per group)

These two experiments further confirmed that glutamate inhibits paravascular movement through the NMDA receptor in a pulsation-dependent manner while GABA has no significant influence on this process.

### The influence of Glutamate and GABA on tracer penetration into parenchyma is AQP4-dependent

Paravascular movement of CSF and ISF is reported to be driven by arterial pulsation and facilitated by an astroglial AQP4 water channel. Given that GABA and glutamate inhibitor accelerated paravascular clearance to significant extents but had distinct effects on dcLN drainage, we propose that AQP4 influences GABA and glutamate activities in different ways.

Initially, AQP4 knockout mice were employed to determine whether GABA and glutamate participate in AQP4-dependent clearance (Fig. 6a). Fluorescence in the paravascular space was slowly increased in AQP4-null mice (Fig. 6b, c), accompanied by little influx into the interstitium (Fig. 6d and e), indicating obvious blockage of para-arterial and interstitium exchange. Glutamate did not influence draining of the tracer into the para-vascular space while APV restored impaired paravascular movement in AQP4-null animals to a remarkable extent (Glutamate vs. vehicle, two-way ANOVA, for interaction factor, *P*=0.9875; APV vs. vehicle, two-way ANOVA, for interaction factor, *P*<0.05) (Fig. 6b). CNQX, GABA, and bicuculline did not alter paravascular movement relative to the vehicle group (CNQX vs. vehicle, two-way ANOVA, for interaction factor, *P*=0.8122; GABA vs. vehicle, two-way ANOVA, for interaction factor, *P*=0.6682; Bicuculline vs. vehicle, two-way ANOVA, for interaction factor, *P*=0.8934) (Fig. 6c). Glutamate almost impaired tracer influx with nearly no penetration of fluorescence into the parenchyma and CNQX failed to facilitate tracer penetration into the parenchyma. APV accelerated paravascular movement but did not promote tracer penetration into the parenchyma (Glutamate vs. vehicle, two-way ANOVA, for interaction factor, *P*=0.6300; CNQX vs. vehicle, two-way ANOVA, for interaction factor, *P*=0.3194; APV vs. vehicle, two-way ANOVA, for interaction factor, *P*=0.5353) (Fig. 6d). Moreover, GABA and bicuculline did not alter the dynamic pattern of the fluorescent signal in parenchyma (GABA vs. vehicle, two-way ANOVA, for interaction factor, *P*=0.4193; Bicuculline vs. vehicle, two-way ANOVA, for interaction factor, *P*=0.2425;) (Fig. 6e).

**Fig. 6.**
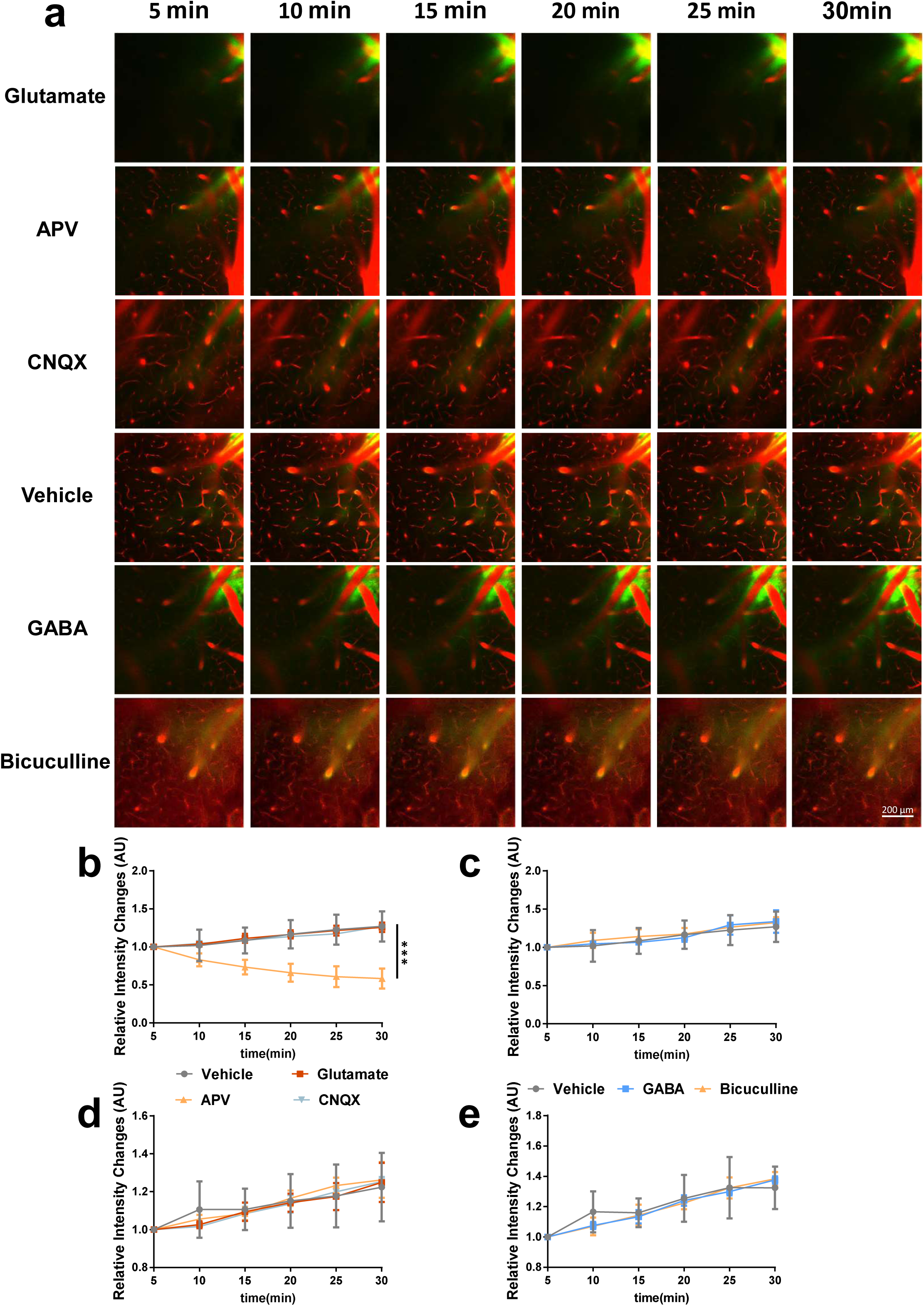
Effects of Glutamate and GABA on tracer penetration into parenchyma are AQP4-dependent. **(a)** *In vivo* two-photon imaging of tracer clearance through the glymphatic pathway after intracisternal injection of GABA/glutamate and the respective antagonists in AQP4^-/-^ mice. **(b, c)** Quantification of CSF tracer influx into the surrounding parenchyma via 3D reconstruction and paravascular CSF tracer clearance at 100 μm below the cortical surface **(d, e)**. APV strongly accelerated paravasular movement while no significant changes were observed in mice receiving glutamate and CNQX. GABA and bicuculline did not affect paravascular movement. No significant changes in tracer penetration into the interstitium were observed among the experimental groups, compared with the vehicle group (n=6 mice per group).

Based on the collective data, we concluded that glutamate inhibited paravascular movement though the NMDA receptor, which was pulsation-dependent, and suppressed tracer penetration into the parenchyma and interstitial fluid drainage though the AMPA/kainate receptor which was AQP4-dependent. GABA did not appear to influence paravascular movement but accelerated glymphatic clearance in an AQP4-depedent manner.

### Hypertension impairs glymphatic clearance of Aβ through reduction of arterial pulsation

Given that glymphatic clearance of interstitial solutes is highly dependent on vascular pulsatility, we hypothesized that hypertension influences pulsatility, in turn, impairing lymphatic clearance. The Ang-II hypertension model was applied to test this theory. After chronic infusion of Ang-II for 28 days, mice displayed a marked increase in systolic blood pressure (SBP) and diastolic blood pressure (DBP), compared to vehicle-infused mice (Fig. 7g, h) (vehicle vs. hypertension, for SBP, *t*=4.429, *P<*0.01, for DBP, *t*=5.326, *P<*0.001). As expected, pulsatility was significantly reduced among surface/penetrating artery and surface/ascending veins (hypertension vs. vehicle, for surface artery, *t*=5.485, *P*<0.001; for penetrating artery, *t*=4.263, *P*<0.01; for surface vein, *t*=12.320, P<0.001; for ascending vein, *t*=5.257, *P*<0.001) (Fig. 7s) in parallel with glymphatic clearance impairment (hypertension vs. vehicle, for paravascular movement, two-way ANOVA, for interaction factor, *P*<0.05; for interstitial movement, two-way ANOVA, for interaction factor, *P*<0.001) (Fig. 7k-m) and obvious enlargement of Virchow-Robin Space (VRS) around the arteries (Fig. 7e). Moreover, we observed no significant differences in GFAP expression and AQP4 polarization (Fig. 7q, t, u). To examine cerebral Aβ deposition *in vivo*, we performed two-photon imaging of fibrillar amyloid plaques pre-labeled with the fluorescent fibrillar amyloid-binding dye, FSB, while simultaneously labeling cerebral vessels with Rhodamine B. *In vivo* imaging revealed significant Aβ deposition on cerebral vessel walls, and to some extent, in surrounding tissues (Fig. 7a-d). This finding was in keeping with the strong staining observed for Aβ 1–40 and weak staining for Aβ 1–42 (Fig. 7p, lower panel). The vehicle group displayed no cerebral FSB-positive amyloid plaques or Aβ staining. Moreover, severe cerebrovascular warp (Fig. 7f) and collagen deposition (vehicle vs. hypertension, for collagen staining, *P*<0.05) (Fig. 7p, r) were detected in hypertensive animals, indicating that hypertension induces substantial changes in vascular structure.

**Fig. 7.**
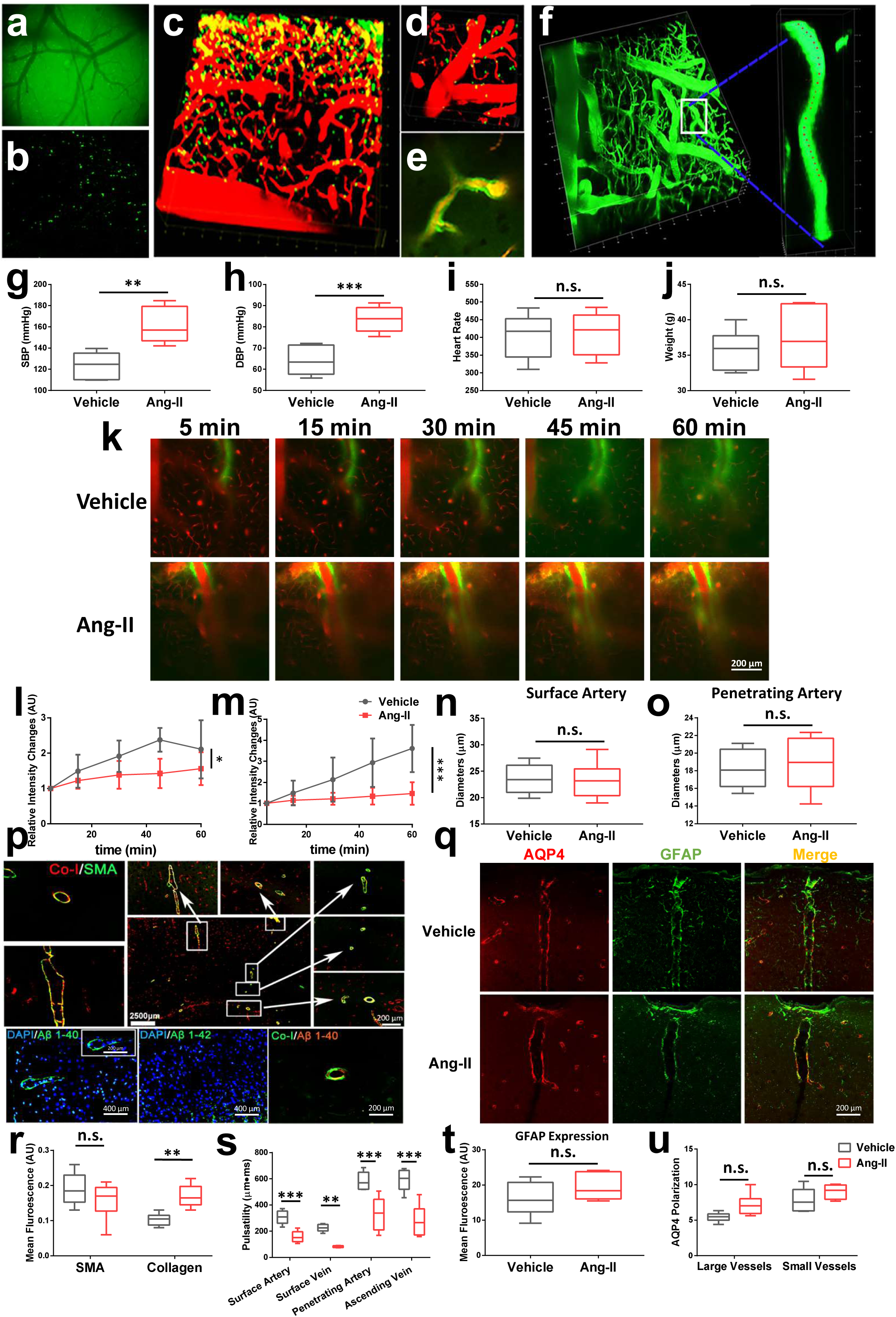
Hypertension induces brain Aβ deposition and glymphatic clearance impairment. **(a–d)** *In vivo* imaging of Aβ deposition in the cerebral cortex. Aβ deposition (FSB, green) was distinct along the vessels. **(e)** Enlarged VRS were observed. **(f)** 3D reconstruction of the vasculature in hypertension mice. The warping vessel is magnified on the right panel. **(g-j)** Ang-II evoked a significant increase in systolic blood pressure (SBP) and diastolic blood pressure (DBP) with no changes in heart rates (HR) and body weight **(k-m).** Glymphatic clearance impairment was evident in hypertension models. **(n-o)** Arterial diameters remained unchanged in hypertension models while vascular pulsatility was severely reduced (n=8-9 vessels per group) **(s)**. **(q)** Representative images of GFAP expression and AQP4 polarization in the cortex. No distinctive changes in AQP4 polarization **(t)** and GFAP expression **(u)** were observed in the hypertension model. **(p, upper panel)** Representative images of smooth muscle actin (SMA) and collagen expression in the cortex. No significant changes in SMA expression **(r)** and greater deposition of collagen **(r)** in vascular walls were observed in hypertension models. **(p, lower panel)** Immunology of Aβ 1–40 and Aβ 1–42 in hypertension model mice. Significant deposition of Aβ 1–40, but not Aβ 1–42, in vessels was observed. **(p, lower panel)** Co-labeling of collagen and Aβ 1–40 in hypertension. Aβ 1–40 co-localized with collagen (n=5-6 mice per group).

### Impairment of glymphatic clearance is associated with Aβ plaque deposition in APP-PS1 mice

Two-photon imaging revealed fibrillar amyloid plaques in the parenchyma, but no evidence of CAA in 7–8-month-old APP-PS1 mice (Fig. 8a), consistent with strongly positive Aβ 1–42 and negative Aβ 1–40 immunofluorescence staining in slices (Fig. 8b). APP-PS1 mice showed significant impairment of glymphatic clearance as indicated by normal paravascular movement but little influx into the interstitium (APP-PS1 vs. WT, two-way ANOVA, for paravascular movement, for interaction factor, *P*=0.2461; for interstitial movement, for interaction factor, *P*<0.001) (Fig. 8d and e), similar to that in AQP4^-/-^ mice. Despite no distinct changes in pulsatility in APP-PS1 mice, increased number of GFAP-positive astrocytes and impaired AQP4 polarization were observed, compared with age-matched WT mice (APP-PS1 vs. WT, for GFAP expression, t=7.527, P<0.001; for AQP4 polarization, among large vessels, *t*=4.215, *P*<0.01; among small vessels, *t*=0.9817, *P*=0.1372) (Fig. 8f-i), indicating that inflammatory responses are activated in the APP-PS1 group that influence the function of AQP4.

**Fig. 8.**
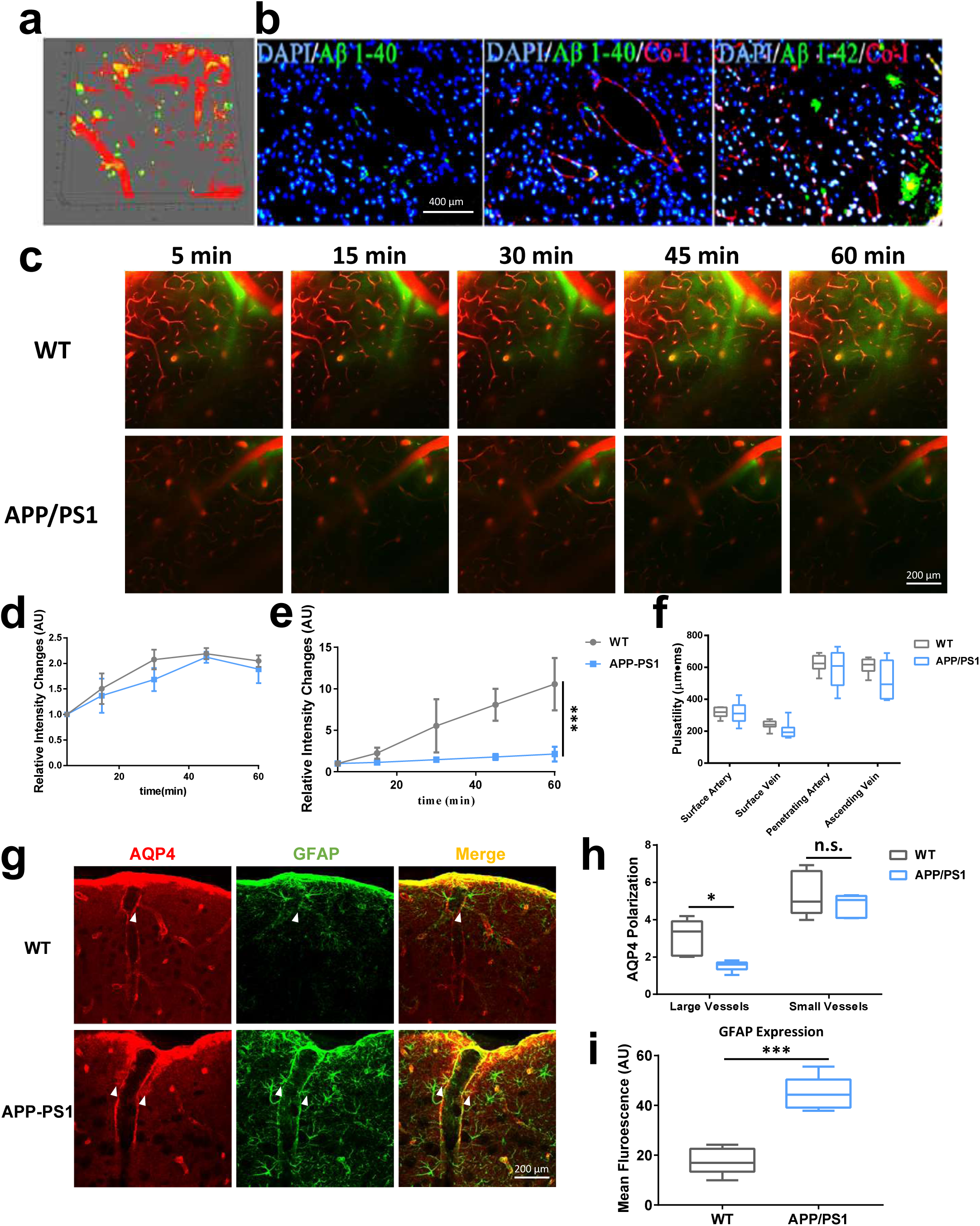
Impairment of glymphatic clearance and deposition of Aβ plaques in APP-PS1 mice. **(a)** *In vivo* imaging of Aβ deposition in the cerebral cortex (FSB: green). Aβ deposition was evident in the parenchyma with no distinct CAA. **(b)** Immunology of Aβ 1–40 and Aβ 1–42 in APP-PS1. Significant numbers of Aβ 1–42-labeled amyloid plaques were observed in the parenchyma, but no marked deposition of Aβ 1–40. **(c)** Representative images of paravascular CSF tracer clearance at 100 μm below the cortical surface in APP-PS1 indicating severe impairment in penetration of fluorescence tracer **(e)** while no changes in paravascular movement was observed **(d). (h-i)** Expression of AQP4 and GFAP in cortex and hippocampus. Compared with WT control mice, APP-PS1 mice displayed significant decrease in AQP4 polarization and exhibited a marked increase in GFAP expression in the cortex (n=6 mice per group). No significant pulsatility changes were observed between APP-PS1 and WT control mice **(f)** (n=7-8 vessels per group).

### Inhibition of glutamate and increase in GABA differentially affect soluble A**β** binding to pre-existing plaques in APP-PS1 and hypertension models

Previously, a method for microinjection of soluble Aβ was developed to assess Aβ binding ability in the brain. Condello and colleagues^20^ showed that Aβ40–555 infusions led to homogeneous binding to amyloid plaques, indicating a direct correlation between soluble Aβ clearance and total binding quantity. We employed this method to examine the roles of glutamate and GABA in glymphatic clearance of soluble Aβ in FAD and hypertension models. Consistently, soluble Aβ microinjected into the parenchyma (Fig. 9b) diffused throughout the hemisphere and rapidly bound to FSB-labeled fibrillar plaques within 10 min (Fig. 9a). After the administration of neurotransmitter into the cisternal, GABA, APV and CNQX inhibited the binding process. Notably, we observed nearly no soluble Aβ binding to FSB-labeled plaques 10 min after administration of GABA, APV and CNQX in both FAD and hypertension models (two-way ANOVA, APP-PS1 vs. GABA, for interaction factor, P<0.001; APP-PS1 vs. APV, for interaction factor, P<0.001; APP-PS1 vs. CNQX, for interaction factor, P<0.001; hypertension vs. GABA, for interaction factor, P<0.001; hypertension vs. APV, for interaction factor, P<0.001; hypertension vs. CNQX, for interaction factor, P<0.001.), supporting the theory that inhibition of glutamate or activation of GABA induces rapid clearance of soluble Aβ (Fig. 9c and d).

**Fig. 9.**
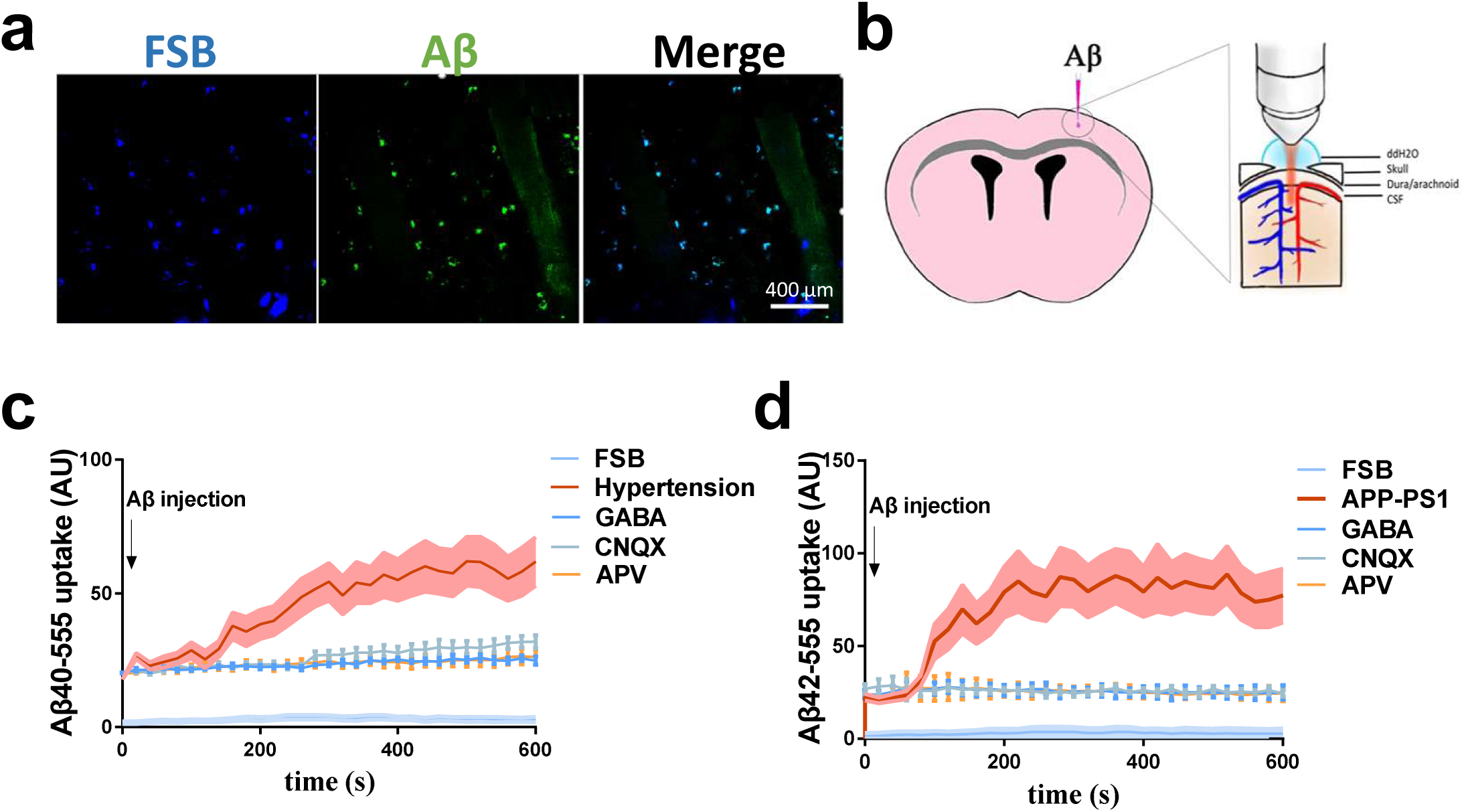
*In vivo* brain imaging of Aβ40–555 binding to existing Aβ deposits. **(a)** High magnification of two-photon images of homogeneous binding of Aβ40–555 (green) to existing amyloid deposits pre-labeled with FSB (blue). **(b)** Schematic diagram of injection of Aβ40–555 into the cortex under a microscope. Quantification of co-labeling of Aβ40-555 with pre-existing amyloid deposits following injection in hypertension **(c)** and APP-PS1 **(d)** mice. Aβ40–555 bound rapidly to pre-existing amyloid deposits and reached a plateau within 5 min (green curve). In the hypertension mouse model **(c)**, APV, CNQX and GABA inhibited binding. However, the CNQX-treated group showed greater binding than the GABA-treated and APV-treated group at 540, 560 and 600 s. In APP-PS1 mice **(d)**, APV, CNQX and GABA inhibited binding with no significant differences (n=5-6 mice per group).

## Discussion

In this study, we have examined the roles of two main neurotransmitters, GABA and glutamate, in modulation of the glymphatic clearance of Aβ. Our experiments suggest that these two neurotransmitters exert opposite effects on glymphatic clearance through distinct pathways. GABA enhanced glymphatic clearance in an AQP4-dependent manner while glutamate suppressed clearance in a manner dependent on arterial pulsation and AQP4 though distinctive receptors.

Arterial pulsation is a main driving force for solute movement in the brain. For example, cerebral arterial pulsation can drive paravascular CSF-interstitial fluid exchange^15^. In addition, arterial pulsation facilitates the exit of ISF solutes in the opposite direction to blood flow along cerebral arteries^21^. In the present study, GABA, APV and CNQX were shown to promote glymphatic clearance. Interestingly, however, APV accelerated paravascular movement, but not GABA or CNQX. In addition, APV promoted clearance of fluorescent tracers in the para-arterial space in AQP4-deleted mice, suggesting that enhancement of glymphatic clearance by APV was not dependent on AQP4 and was more likely acts on blood vessels. Indeed, glutamate is reported to constrict pial arterioles whereas inhibition of glutamate by an ionotropic receptor antagonist dilates arterioles^18^. Because APV and CNQX are both ionotropic receptor antagonists, the issue of why APV but not CNQX promotes paravascular movement of fluorescent tracers was unclear. One possible explanation was that endothelial cells expressed NMDA but not AMPA/kainate. Activation of endothelial NMDA could increase Ca^2+^ influx, which, in turn, reduced pulsatility.

The involvement of AQP4 in solute movement in the brain has been well established^4, 6, 22^. Consistent with previous observations, significant reduction of interstitial clearance was evident in AQP4-/- mice. CNQX-mediated interstitial clearance was proposed to rely heavily on AQP4, in view of the finding that increase in glymphatic clearance was abolished in AQP4-/- mice. Although astrocytes expressed ionotropic receptors, including AMPA/kainate and NMDA, only AMPA was widely expressed in many brain regions and has been confirmed to act as a fully functional receptor. The AMPA receptor functions through mediation of calcium and sodium influx in astrocytic processes^23^. However, the mechanism underlying AQP4 involvement in CNQX-mediated clearance is currently unclear. This process may be mediated indirectly because no direct interactions have been detected between AMPA and AQP4. Glutamate exerts depolarizing effects on astrocytes through ion influx. Upon depolarization of astrocytes, molecules such as AQP4 that normally localize in endfoot membranes are redistributed to non-endfoot membranes, thereby compromising AQP4 function^24^. By inhibiting ion influx, CNQX may therefore counteract glutamate-mediated astroglial depolarization to maintain AQP4 in perivascular endfeet.

Unlike glutamate, GABA had no effect on pulsation. Inhibition of GABA by the GABA_-A_ receptor antagonist, bicuculline, led to a considerable decrease in glymphatic clearance. GABA enhanced glymphatic clearance in wild-type but not AQP4^-/-^ mice, indicating that GABA-mediated clearance was AQP4-dependent. Earlier studies suggested that the GABA receptor co-localizes with AQP4^16, 25^. Further research is warranted to ascertain whether direct interactions between GABA and AQP4 promote glymphatic clearance.

Failure to clear Aβ deposits is linked to a number of Aβ-associated diseases, such as FAD and hypertension^8, 26, 27^. However, brain distribution of Aβ is variable among different diseases^28^. For example, Aβ mainly localizes in brain parenchyma in AD but is deposited along arteries in hypertension^1, 3, 19^. These findings highlighted the complexity of the mechanisms underlying failure of Aβ deposit clearance. Accordingly, we further explored the potential roles of glutamate and GABA neurotransmitters in clearance of Aβ deposits in FAD and hypertension.

Transgenic APP/PS1 mice, a commonly used model of AD, can generate substantial amounts of β-amyloid. Consistent with previous observations, significant levels of Aβ were deposited in the parenchyma in APP/PS1 mice. Similar to AQP4-/- mice, APP-PS1 mice exhibited marked impairment of glymphatic clearance, specifically, an increase in paravascular fluorescence but little influx into the interstitium. While no significant alterations in pulsatility were observed, AQP4 polarization was remarkably impaired and GFAP expression increased in APP-PS1 mice. These findings may explain the impaired interstitium clearance in APP-PS1 mice and shed further light on the mechanisms underlying Aβ deposition in Alzheimer’s disease patients.

Hypertension is a strong risk factor for age-related dementia that is characterized by both beta amyloid (Aβ) deposition and vascular dysfunction^3^. Consistently, Aβ deposition was evident in Ang-□ induced hypertensive animals as well as glymphatic clearance impairment. In parallel, pulsatility was significantly decreased whereas the GFAP and AQP4 levels remained unchanged. Thus, the hypertension-induced decrease in arterial pulsation was possibly responsible for Aβ deposition. Interestingly, GABA, APV and CNQX promoted rapid clearance of soluble Aβ in both FAD and hypertension models, supporting the theory that modulation of neurotransmitters may serve as an effective therapeutic strategy for removal of Aβ deposits in the brain. However, it should be noted that CNQX was less effective than GABA in soluble Aβ clearance in the hypertension model, which may be attributable to differences in their mechanisms of action. Unlike GABA, CNQX does not interact directly with AQP4.

Our study had a number of limitations that need to be addressed. Previous reports suggest that Aβ can be removed from the brain via various clearance systems (most importantly, the blood–brain barrier (BBB)). As the glymphatic pathway flushes Aβ towards the perivascular space, which may influence clearance through BBB^1, 15, 28^, the issue of whether GABA and glutamate modulate BBB permeability or act via interactions with receptors for advanced glycation end products (RAGE) or low density receptor-related protein-1 (LRP1) requires further investigation.

In conclusion, the neurotransmitters glutamate and GABA exerted distinct modulatory effects on glymphatic clearance. The collective data from our study presented novel insights into the mechanisms underlying Aβ drainage that may be effectively applied for treatment of Alzheimer’s disease and hypertension.

## Acknowledgments

This work was supported by grants from the National Natural Science Foundation of China (Grant Numbers: 81572224, 81572230, 81772438, and 81671102), The National Key Research and Development Program of China, Stem Cell and Translational Research (2017YFA0105104), Science and Technology Planning Project of Guangdong Province, China (Grant Numbers: 2016A020213003, 2016B030230002, and 2017A040406007), Science and Technology Planning Project of Guangzhou, China (Grant Number: 2016201604030036), and the National Key Clinical Department, National Key Discipline, Guangdong Provincial Key Laboratory for Diagnosis and Treatment of Major Neurological Disease (Grant Numbers: 2014B030301035).

## Author contributions

Yi-wei Feng, Qun Zhang, Xiao-fei He, Dong-xu Liu, Dan Wu, Ge Li, Cheng Wu and Feng-yin Liang performed the experiments. Yi-wei Feng, Qun Zhang and Xiao-fei He drafted the manuscript. Guang-qing Xu, Yue Lan and Zhong Pei conceived and designed the research. Zhong Pei and Guang-qing Xu edited and revised the manuscript. Guang-qing Xu, Yue Lan and Zhong Pei approved the final version of the manuscript.

